# Maximizing viral detection with SIV droplet digital PCR (ddPCR) assays

**DOI:** 10.1101/668285

**Authors:** Samuel Long, Brian Berkemeire

## Abstract

Highly sensitive detection of HIV-1 nucleic acids is of critical importance for evaluating treatment interventions superimposed on combination antiretroviral therapy (cART) in HIV-1 infected individuals. SIV infection of rhesus macaques models many key aspects of human HIV-1 infection and plays a key role in evaluation of approaches for prevention, treatment and attempted eradication of HIV infection. Here we describe ultrasensitive droplet digital PCR (ddPCR) DNA and RT-ddPCR RNA assays for detecting simian immunodeficiency virus (SIV) on the Raindance ddPCR platform. We demonstrate that RainDance ddPCR can tolerate significantly higher cell DNA input without inhibition on a per reaction basis, exceeding the Bio-Rad ddPCR per-reaction input limit by about 18-fold, effectively increasing viral detection sensitivity by allowing a large quantity of sample to be analyzed in each reaction. In addition, the combination of a high processivity RT enzyme and RainDance ddPCR could overcome inhibition in severely inhibited viral RNA samples. These assays offer valuable tools for assessing low level viral production/replication and strategies for targeting residual virus in the setting of cART suppression of viral replication. The methodologies presented here can be adapted for a broad range of applications where highly sensitive nucleic acid detection is required.

## Introduction

The ability to accurately detect and quantitate HIV-1 nucleic acid levels is important for evaluating the efficacy of antiretroviral drug therapies and monitoring disease status in HIV-1 infected patients. Combination antiretroviral therapy (cART) has resulted in suppression of viral load in patients to levels that require the use of ultrasensitive (e.g. single copy sensitivity) assays to monitor patient response and make therapeutic decisions (Palmer et al 2003). Viral detection and quantitation can however be complicated by technical issues related to sample inhibition from two sources. The nucleic acid input required for ultralow level viral detection can significantly exceed the per reaction nucleic acid input capacity of real time PCR or a Bio-Rad droplet digital (ddPCR) instrument platforms. Exceeding the input capacity of these platforms can lead to significant reaction inhibition during quantification (Strain et al, 2013). Another factor that often confounds viral nucleic acid quantification is inhibitors that are introduced during sample procurement or co-purify with specimen-derived nucleic acids during extraction. Inhibition can occur either at the reverse transcription stage (in the case of RNA samples), or the PCR amplification/quantification stage (such as in qPCR, for cDNA and DNA samples), or both. Anticoagulants such as heparin are known to inhibit various steps of nucleic acid quantification, and can be avoided at the sample procurement step. Other potential inhibitors, either inherent in the source specimens, or introduced during extraction, can also inhibit nucleic acid quantification steps and are difficult to remove. These two sources, individually or in combination, can lead to lowered sensitivity of viral detection (even in cases where an optimally sensitive qPCR assay is used) and confound the interpretation of viral load results.

Droplet digital PCR (ddPCR) has seen increased utilization in both nucleic acid quantification and next generation sequencing. ddPCR platforms such as RainDance achieve micro-partitioning of analyte by emulsification of the aqueous PCR reaction mixture into picoliter droplets of thermostable oil. For quantification applications, these platforms measure the end-point fluorescence of a large number of droplets of the same sample (for RainDance, up to 10 million droplets per reaction), prepared in the limiting dilution range, instead of measuring the real-time increase of fluorescence intensity within one sample. Quantification is achieved using Poisson statistics by counting fluorescence-positive and total droplet numbers. Compared to real time PCR methods, ddPCR, which is based on a simple endpoint PCR digital positive/negative readout, avoids the need for assay calibration standards, can potentially alleviate assay competition in multiplex assays, and has less stringent requirements for primer/probe sequence match to target sequence. ddPCR can provide greater precision at low analyte copy numbers compared to the 1.25-1.5 fold minimal difference demonstrable under ideal conditions with real-time PCR (Weaver et al.), and the sensitivity for detecting rare alleles can be much higher than that for real time PCR (which is typically 1-10%).

Besides the general advantages that apply to all ddPCR platforms, such as being less prone to inhibition and having better data reproducibility, especially at low copy numbers, the RainDance platform has a few unique advantages: (1) It is an open platform which allows use and optimization of a variety of assays, reagents, and lab-developed protocols. This enables potential migration of assays developed using non-ddPCR conditions to the ddPCR platform; (2) The 10 million droplet capacity per reaction offers a wider dynamic range than other existing ddPCR platforms, allowing analysis of a greater range of sample concentrations for any given precision; (3) The platform allows multiplexing of more than 2 assays by varying probe color or intensity for different targets (Zhong et al.).

We report the development, validation and optimization of an ultrasensitive RainDance ddPCR DNA assay, and an ultrasensitive RainDance reverse transcription (RT)-ddPCR RNA assay for simian immunodeficiency virus (SIV), which is widely used in nonhuman primate models for HIV/AIDS studies. We investigate the feasibility of utilizing the RainDance ddPCR platform to overcome viral detection inhibition caused by high nucleic acid input (for proviral detection), or caused by inhibitor(s) that copurify with specimen-derived nucleic acids. We demonstrate that RainDance ddPCR can tolerate significantly more cell DNA input (without inhibition) on a per reaction basis, exceeding the Bio-Rad per-reaction input limit (Strain et al. 2013) by about 18-fold. In addition, the RainDance platform minimizes the impact of copurifying inhibitors, allowing quantification in RNA sample(s) that are not quantifiable by real time PCR.

Rhesus macaques infected with SIV are the animal model of choice for many studies of key aspects of HIV biology, including characterization of viral reservoirs during suppressive cART, and evaluation of approaches to target this residual virus, applications requiring high sensitivity for viral quantification. Even with optimized assays, maximal sensitivity necessitates a large amount of input nucleic acid. The capacity of the RainDance platform to overcome inhibition during nucleic acid quantification can have broad applications in other areas as well where large quantities of input DNA is required, or quantification of inhibited/challenging RNA samples is involved.

## Results

### Significant inhibition in viral quantification at high nucleic acid input levels on a real time PCR platform

During real-time PCR analysis of SIV infected Rhesus macaque tissue-derived DNA for SIV DNA levels it is not uncommon to observed inhibition when performing viral quantitation from typical snap frozen tissue specimens. In the set of tissue DNA samples shown in Table 1, we measured copy number for SIV gag DNA and for a single copy genomic sequence from CCR5 with real-time PCR assays (i.e. the CCR5 copy number in uninhibited reactions (“1:10 diluted”) serve as an indicator of total DNA input in each reaction). We spiked 2 of 12 reactions with 1,000 copies of a SIV positive control template, and determined copy numbers for the other 10 replicate reactions. As shown in the undiluted sample portion of the table, reactions were markedly inhibited across all tissues, with 1,000 copies of spiked SIV DNA registering on average at less than 1 copy in the background of the extracted tissue DNA. When the samples were diluted 1:10 and re-tested, the spiked samples showed appropriate copy numbers of ∼ 1,000, indicating that the inhibition had been relieved through dilution. For the 1:10 diluted samples, CCR5 copy numbers ranged from ∼5 × 10^E5 to ∼9 × 10^E5, corresponding to approximately 2.5-4 × 10^E6 diploid genome cell equivalents (2 copies of the amplified CCR5 sequence/diploid cell) DNA input per reaction for undiluted samples. Comparison of results for SIV DNA in the diluted vs. undiluted samples underestimated actual copy numbers by up to 1,000 fold; quantitation of CCR5 copy numbers was less inhibited in the undiluted samples (compare CCR5 average copies (single stranded) column in “1:10 diluted” vs. “Undiluted”), presumably due to greater template abundance. Together, these results indicate that input DNA level at 2.55 × 10^E6 cells equivalent (based on the rectum sample, which has the lowest CCR5 average copy number in 1:10 diluted sample) or above in each “undiluted” reaction, alone or in combination with inhibitors in the extracted nucleic acid can lead to significant PCR inhibition on the real time PCR platform to compromise assay accuracy.

**Table 1.**
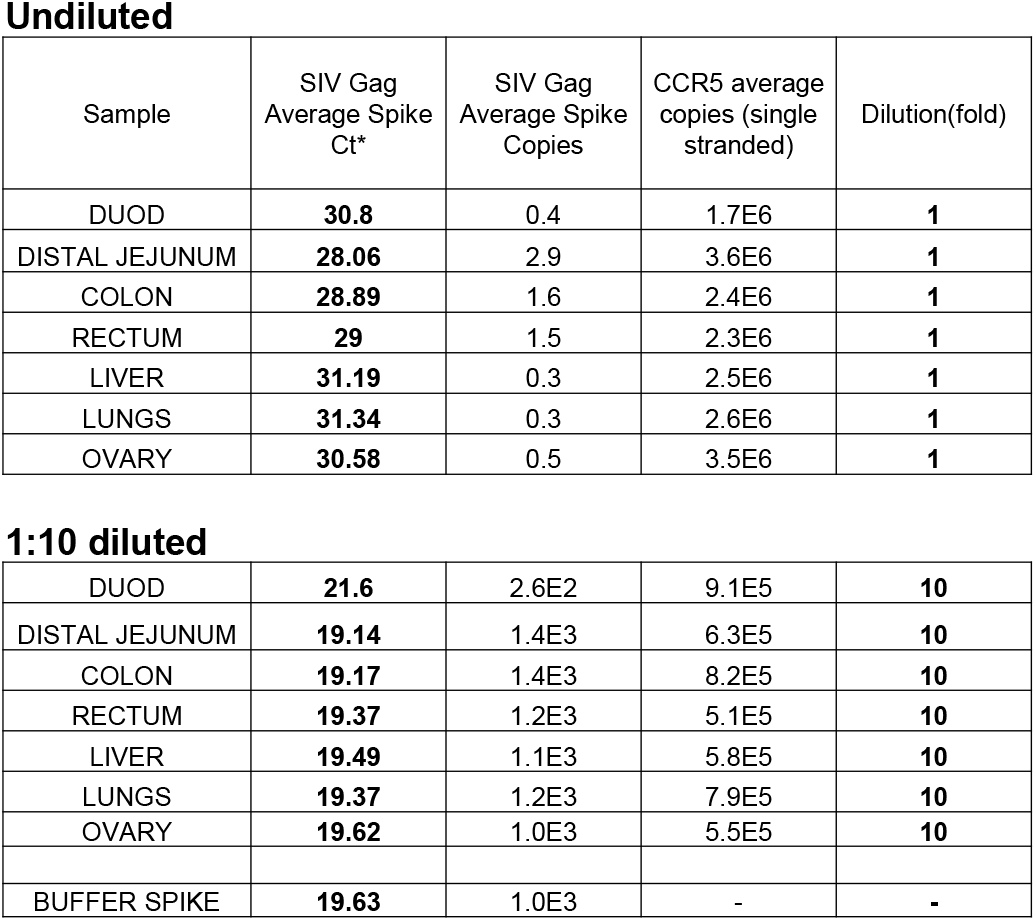
Tissue DNA inhibition for SIV viral quantification. DNA was extracted from 7 tissues from a necropsied Rhesus macaque (310-08) that was infected with SIVmac239 (see Materials and Methods section for details). DNA samples (undiluted or 1:10 diluted) were first subject to preamplification, and 10% of each preamp product was subject to duplex qPCR of Rhesus macaque SIV gag and CCR5 (Hansen et al 2013). 1000 copies of SIV DNA standard was spiked into 2 reactions containing the preamp product to assess PCR inhibition in tissue DNA, while a buffer spike (1000 copies of SIV DNA standard spiked into buffer) served as “no inhibition” control. Average spike copies for SIV gag was derived from average spike Ct (compared to buffer spike Ct), while CCR5 average copies were derived from a CCR5 standard curve. SIV copy number was derived directly with reference to an SIV standard curve.

| Sample | SIV Gag<br>Average Spike<br>Ct* | SIV Gag<br>Average Spike<br>Copies | CCR5 average<br>copies (single<br>stranded) | Dilution(fold) |
| --- | --- | --- | --- | --- |
| DUOD | 30.8 | 0.4 | 1.7E6 | 1 |
| DISTAL JEJUNUM | 28.06 | 2.9 | 3.6E6 | 1 |
| COLON | 28.89 | 1.6 | 2.4E6 | 1 |
| RECTUM | 29 | 1.5 | 2.3E6 | 1 |
| LIVER | 31.19 | 0.3 | 2.5E6 | 1 |
| LUNGS | 31.34 | 0.3 | 2.6E6 | 1 |
| OVARY | 30.58 | 0.5 | 3.5E6 | 1 |
| 1:10 diluted |  |  |  |  |
| DUOD | 21.6 | 2.6E2 | 9.1E5 | 10 |
| DISTAL JEJUNUM | 19.14 | 1.4E3 | 6.3E5 | 10 |
| COLON | 19.17 | 1.4E3 | 8.2E5 | 10 |
| RECTUM | 19.37 | 1.2E3 | 5.1E5 | 10 |
| LIVER | 19.49 | 1.1E3 | 5.8E5 | 10 |
| LUNGS | 19.37 | 1.2E3 | 7.9E5 | 10 |
| OVARY | 19.62 | 1.0E3 | 5.5E5 | 10 |
| BUFFER SPIKE | 19.63 | 1.0E3 | - | - |

### qPCR to ddPCR migration based on an existing ultrasensitive SIV gag DNA assay

As digital droplet PCR (ddPCR) platforms are less prone to inhibition, we tested whether an ultrasensitive SIV gag DNA assay (Li et al, 2016) in its existing form (i.e. primer and probe sequences, and master mix condition) could be migrated onto the RainDance ddPCR platform. The main issue encountered during this migration attempt was insufficient cluster separation (i.e. separation of PCR negative droplets from positive, SIV DNA-containing droplets) which can lead to low signal to noise ratio (Figure 1A), when testing either extracted tissue DNA containing preamplified SIV DNA, or reactions spiked with non-preamplified SIV DNA standards as input template. (In all ensuing ddPCR experiments, non-preamplified tissue DNA or SIV DNA standard spike controls were used.) To overcome this issue, we altered several parameters in the master mix used in the real time assay format, including MgCl_2_ concentration (Figure 1B), primer concentration, probe concentration, PCR annealing temperature and DNA polymerase concentration (data not shown). Among these modifications, changing MgCl_2_ concentration proved to have the most dramatic effect, with increasing MgCl_2_ concentration leading to better cluster separation.

**Figure 1.**
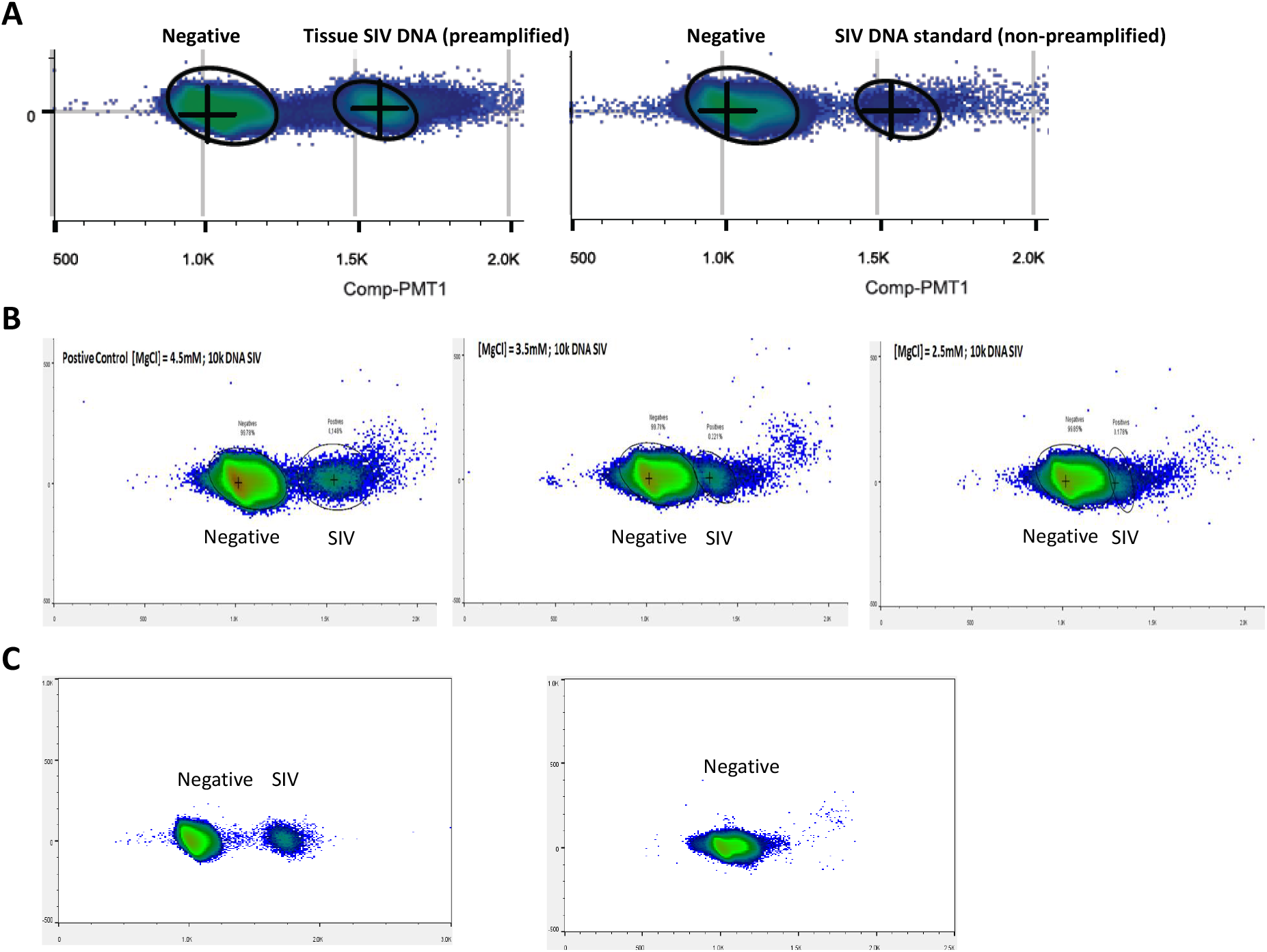
ddPCR optimization based on existing SIV gag DNA real time qPCR assay and condition. (A) Direct migration of the qPCR SIV gag DNA assay in existing format onto Raindance ddPCR platform. Left: tissue DNA containing preamplified SIV DNA as template; Right: SIV DNA standard spike-in in buffer as template. (B) The effect of modifying MgCl_2_ concentration on cluster separation. (C) Background issue in the SIV target detection region using existing assay’s master mix with optimized MgCl_2_ concentration.

The most optimal condition for maximal cluster separation based on the existing master mix condition was achieved by increasing the MgCl_2_ concentration to 5.5 mM, while keeping all other conditions unchanged (Figure 1C). However, the main disadvantage associated with this modified condition was significant background in the target detection region when no SIV DNA template was present (Figure 1C). This limited the utility of simply transferring the existing real time assay conditions but with a higher MgCl_2_ concentration for ultralow level SIV viral detection on the ddPCR platform.

### Optimization of ddPCR assay conditions

To achieve improved cluster separation and clean background (i.e. signal to noise ratio) in the target detection region on the RainDance ddPCR platform, we introduced the following changes to the assay format and/or master mix condition: (1) For PCR cycling, the manufacturer-recommended 45 cycles were reduced to 40. This was based on the estimate that in target-containing droplets, dNTP and probe will be exhausted at lower than 40 cycles (assuming 100% PCR efficiency; this may overestimate how quickly the reagents will be depleted). Reducing PCR cycles is expected to improve cluster separation, as the 5 additional PCR cycles may lead to increased background signal intensity and consequently less optimal cluster separation, with negligible incremental sensitivity. (2) The existing assay’s probe contains a single blackhole quencher (BHQ). We considered switching the probe to a double-quenched probe or TaqMan MGB probe. In doubly quenched probes (Integrated DNA Technologies) there is an additional ZEN quencher 9 base pairs from the dye in addition to the Iowa Black quencher. Dually quenched negative droplets are expected to provide increased signal-to-noise ratio allowing for better cluster separation. TaqMan MGB probes use a non-fluorescent quencher (NFQ) that offers low background signal, although a direct comparison of the background signals between TaqMan MGB probes and probes containing a single blackhole quencher has not been described. (3) Several PCR mastermixes were tested, including TaqMan universal mastermix (Life Technologies), Quanta Toughmix (Quantabio), TaqMan genotyping mastermix (Life Technologies) and Quanta genotyping mastermix (Quantoabio) (the last two mastermixes are optimized for end point PCR).

In evaluating assay and master mix performance, in addition to cluster separation and background signal, we also considered signal count (i.e. whether the positive droplet count agrees with the number of input molecules) and signal cluster diffuseness. Among all the conditions tested, an MGB probe combined with TaqMan genotyping master mix yielded the best cluster separation, signal count, least cluster diffusion and cleanest background (Figure 2A, B), either when positive control template-spiked samples were used or tissue DNA containing SIV was used, while other combinations suffered from deficiencies in one or more criteria. For example, for a doubly quenched probe, Quanta ToughMix and TaqMan Genotyping Mastermix both had sufficient cluster separation and signal counts, but there was low level background in target signal region for SIV DNA negative samples under both conditions (Supplementary Figure 1A). Quanta ToughMix in combination with MGB probe performed adequately with spiked template samples (Supplementary Figure 1B) with somewhat diffuse target signal clusters. However, the Quanta ToughMix performed poorly when tested with tissue DNA as input (Supplementary Figure 1C; i.e. low background in target signal region, and diffuse cluster). A summary of all conditions (including different probe, master mix and input sample combinations) tested and the observed issue(s) associated with each condition is shown in Supplementary Figure 1D.

**Figure 2.**
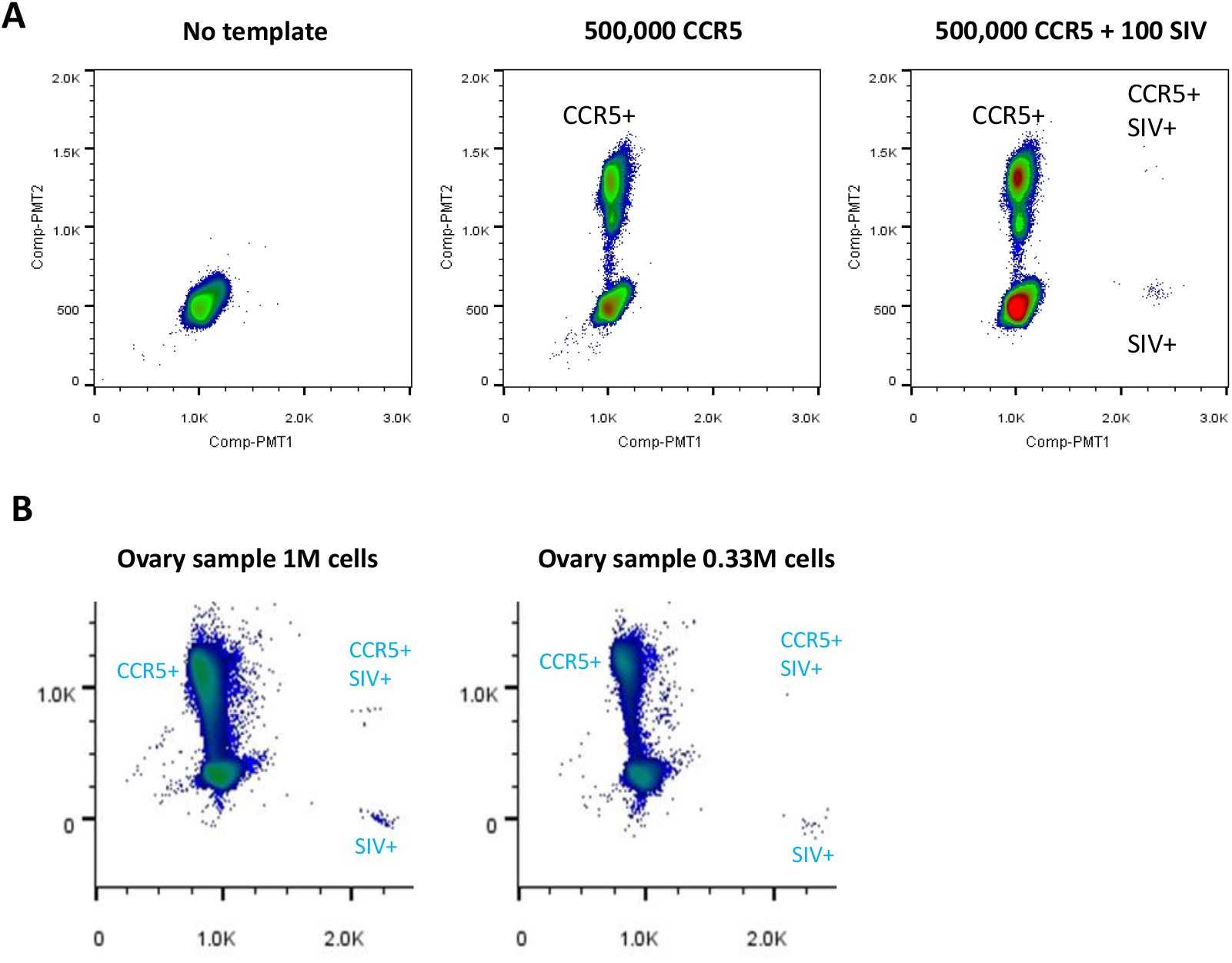
Identifying an optimal ddPCR assay format and condition. Performance of an MGB probe-based SIV gag ddPCR assay using TaqMan genotyping mastermix on (A) SIV and CCR5 spike-in templates and (B) ovary tissue DNA from a Rhesus macaque (311-08) infected with SIVmac239.

### SIV ddPCR DNA assay performance and characterization

We further characterized the MGB probe-based SIV ddPCR DNA assay (in TaqMan genotyping master mix)’s input tolerance to DNA and the assay’s detection sensitivity. DNA input tolerance was tested by (1) visually observing droplet integrity on the RainDance Source instrument as the droplets were being generated in each lane and moved through the device, and by (2) Source machine counts of total intact droplet number for each sample input level (in increments of 1 million cell equivalent of genomic DNA per lane) after dropletization. It was previously shown that sample input tolerance for the Bio-Rad ddPCR platform was below 3 ug per reaction (Strain et al, 2013), i.e. droplet deformation and number decrease start at 3 ug DNA input per well (equivalent to less than a half million mammalian cell equivalents) on the Bio-Rad platform. We observed that the RainDance instrument can tolerate at least 52.8 ug DNA (8 million Rhesus macaque cell equivalents of DNA input) per reaction without compromising droplet integrity or quantity (Figure 3A; Supplementary Table 1). This input tolerance is significantly higher (∼35 fold) than the Bio-Rad platform, even when equal reaction volumes are compared (RainDance vs. Bio-Rad reaction volume: 50 uL vs. 20 uL). As 52.8 ug per reaction was the highest input amount tested, without evidence of compromise of droplet formation, it is possible that the RainDance platform can tolerate even higher levels of nucleic acid on a per reaction basis at the droplet formation step.

**Figure 3.**
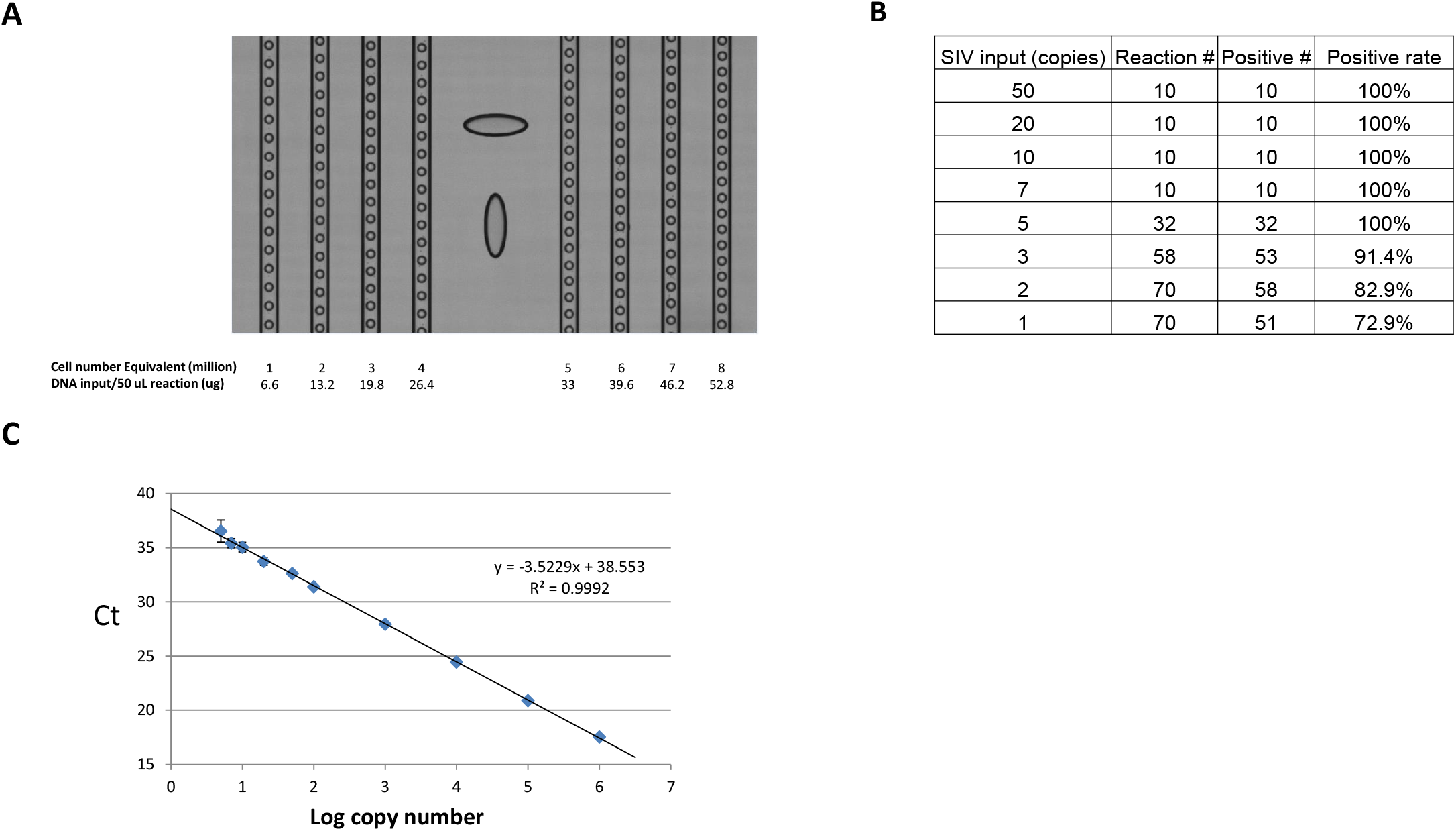
SIV ddPCR DNA assay characterization. (A) Sample DNA input tolerance at the droplet formation step. Droplet integrity was monitored by examining a portion of the droplets in each lane as they moved through the Source instrument during dropletization. In addition, total droplet number for each input level after dropletization (retrieved from the “RainDrop Run Completion” screen) served as another indicator of sample DNA input tolerance (Supplementary table 1). DNA sample used was duodenum DNA from Rhesus macaque 313-08. (B) Estimation of the limit of detection (LoD) of the ddPCR assay based on the Digital MIQE Guidelines (Huggett et al). According to the guidelines, when running costs preclude optimization using ddPCR, qPCR can be used to determine certain assay parameters. (C) Performance of the SIV ddPCR assay in TaqMan genotyping mastermix in qPCR format. SIV gag DNA standard was diluted with buffer diluent. The standards were assayed as described in Materials and Methods in the following replicate format: 1 million down to 100 copies input per reaction: each in triplicates; 50, 20, 10, 7 and 5 copies input per reaction: each in 10 replicates. The data were plotted and analyzed according to the routine analyses provided in the software package with the ABI 7500 SDS instrument.

The MGB probe-based SIV gag ddPCR DNA assay has an estimated LoD of 4 copies per reaction in qPCR format using the Taqman genotyping master mix (Figure 3B), making it close to a single copy assay (i.e. an LoD of 3 copies per reaction, giving a 95% probability of detecting a single copy if three replicates are tested). Due to the potential high cost of repeating the LoD experiment (Figure 3B) on the RainDance ddPCR platform, we followed the Digital MIQE guidelines (Huggett et al), and obtained an estimation of the assay’s LoD based both on the qPCR data and on the ddPCR platform by using a lower number of ddPCR replicates and required the false negative rate to be below 5% (i.e. all 10 replicates have to be positive). All 10 reactions with 3 copy SIV DNA input were positive (Supplementary table 2).

As the main intended utility of the SIV ddPCR assay was to detect and/or quantify ultralow level SIV virus, we confirmed that the assay can reliably detect single digit levels of SIV viral DNA in samples consisting of either spiked-in SIV DNA or genomic DNA extracted from tissues from SIV-infected Rhesus macaques chronically suppressed with cART (Figure 4, SIV DNA standards spiked in 1 million uninfected Rhesus macaque cell equivalent genomic DNA; 2B and 2B, tissue DNA). For example, in the tissue DNA shown in Figure 2B, 1 doubly occupied (SIV + CCR5) droplet and 15 singly occupied (SIV) droplets were detected. These results suggest that the assay is suitable for direct analysis of tissue-derived nucleic acid samples containing low level SIV DNA, without an enrichment step such as a preamplification PCR reaction. The linear dynamic range of the assay was confirmed to be at least up to 1 million copies (test upper limit) of viral nucleic acid per reaction (Supplementary Figure 2).

**Figure 4.**
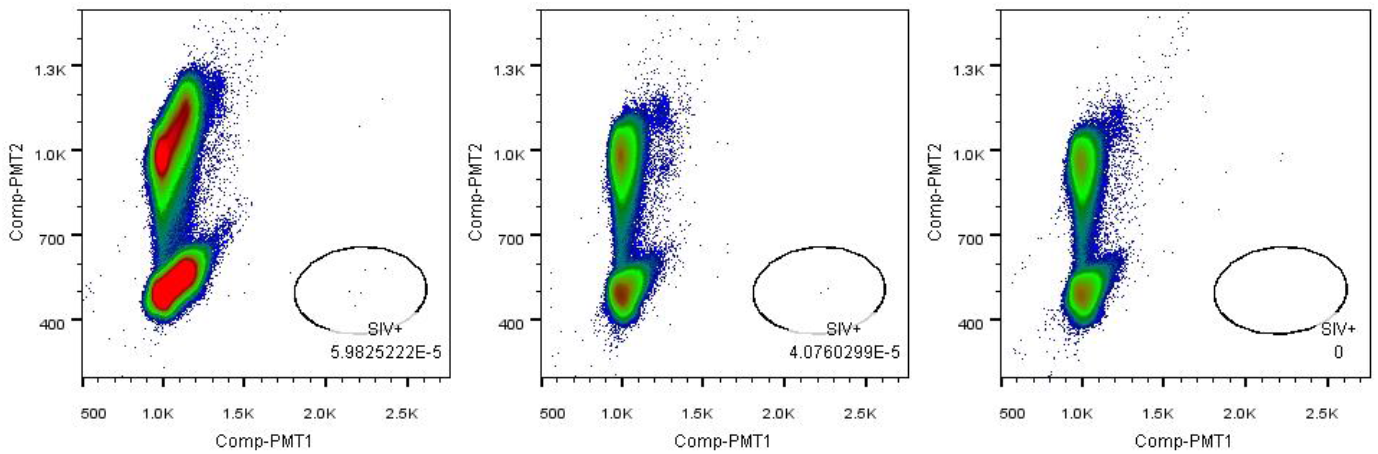
SIV ddPCR DNA assay performance. SIV ddPCR DNA assay detection of low single digit level SIV DNA input. 3 (left), 2 (middle) or 0 (right, negative control) copies of SIV DNA standard were spiked in 1 million Rhesus macaque PBMC equivalent of genomic DNA background each. Measured SIV DNA spike-in amount: 5 (left), 2(middle), 0(right).

### Comparison between ddPCR and qPCR quantitation results in low viral DNA range

To compare the performance between the new ddPCR assay (with MGB probe and TaqMan genotyping mastermix) and a non-preamplifed version of the existing real time SIV gag qPCR assay (Li et al 2016) especially in low viral DNA range, DNA extracted from 1 million PBMC derived from an SIVmac239 infected Rhesus macaque receiving suppressive cART was subjected to either direct analysis by ddPCR (Table 2, format 1), or analyzed in 10 qPCR reactions (format 2; each containing non-preamplified, 0.1 million cell equivalent DNA) (viral load deduced from Poisson statistics). Similar results were achieved between the 2 analyses. In addition, when 2 million cell equivalent DNA was divided among 40 qPCR reactions (format 3) and viral load per million cells was deduced from Poisson statistics, a similar result was obtained. We conclude that the ddPCR-derived (format 1) and Poisson-based qPCR-derived (formats 2 and 3) viral loads were consistent with each other at low viral DNA range.

**Table 2.** Comparison between ddPCR and dPCR/qPCR hybrid assay format for viral quantification. DNA extracted from 1 million PBMC derived from an SIVmac239-infected Rhesus macaque was subjected to either analysis by ddPCR (format 1), or analyzed in 10 qPCR reactions (format 2; each containing 0.1 million cell equivalent DNA) (viral load deduced from Poisson adjustment). In addition, 2 million cell equivalent DNA from the same animal was divided among 40 qPCR reactions (format 3) and viral load per million cells deduced from Poisson statistics. For both ddPCR and qPCR, PBMC DNA from naïve (uninfected) Rhesus macaque was used as negative control.

| **Format** | **Input** | **SIV DNA/million cells** | **Data analysis** |
| --- | --- | --- | --- |
| 1. ddPCR | 1M PBMCs | 17 | Count + Poisson |
| 2. qPCR | 0.1M cells/rxn * 10 rxns | 23 | Poisson (9/10 positive) |
| 3. qPCR | 0.05M cells/rxn * 40 rxns | 21 | Poisson (26/40 positive) |

### SIV ddPCR DNA assay overcomes PCR inhibition due to high input nucleic acid levels

Next, we investigated the utility of the SIV ddPCR DNA assay to quantify the SIV DNA viral load in tissue DNA samples that showed significant inhibition during qPCR analysis (Figure 5A left), and compared the results to qPCR results for 1:10 diluted sample. We observed that with up to 4 million cell-equivalents of DNA input (26.4 ug) per reaction, there was no detectable inhibition in ddPCR quantification of SIV DNA viral load, while at a similar genomic DNA input level, qPCR showed ∼99% inhibition in SIV DNA quantification for undiluted DNA sample. In comparison, Strain et al reported that loading of >1.5 ug of DNA per Bio-Rad ddPCR reaction greatly reduced the measured copy number of target sequences for all samples tested, suggesting significant inhibition at or above 1.5 ug sample input on that platform. This result indicates that each RainDance ddPCR reaction can tolerate significantly more DNA input both at the droplet formation step (compared to Bio-Rad ddPCR platform), and at the chemical reaction level (compared to both Bio-Rad ddPCR and qPCR).

**Figure 5:**
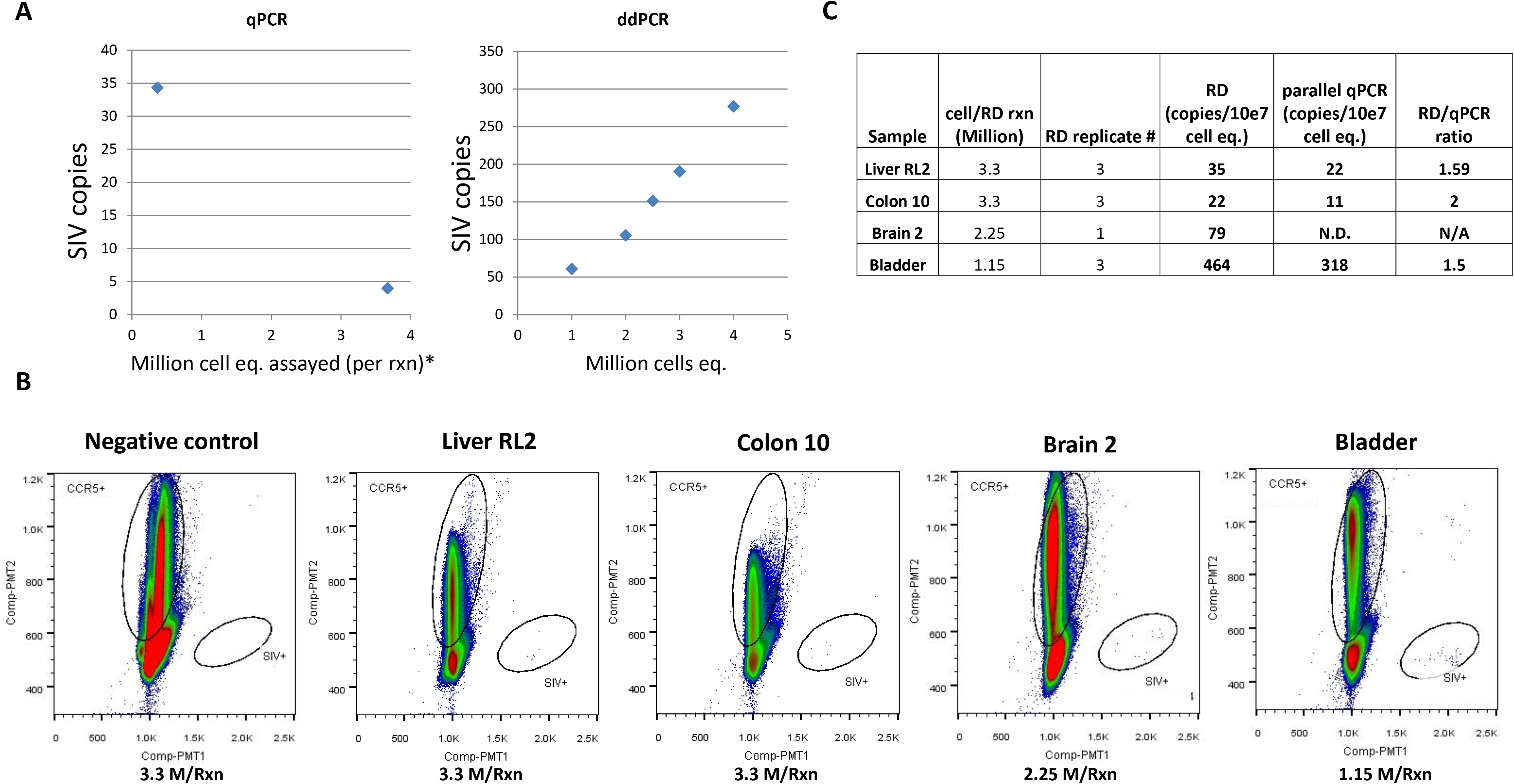
ddPCR and qPCR comparison. (A) Sample inhibition comparison between ddPCR and qPCR. An ovarian DNA sample (Rhesus macaque 313-08) was subjected to nested (i.e. preamplified) qPCR analysis or direct RainDance (“RD” in the table header) ddPCR analysis for SIV DNA viral load. At 3.7 M cell input per reaction (at the preamp step), SIV qPCR signal is inhibited by 99%, while at 4 M cell input per direct ddPCR reaction, SIV signal is not inhibited. (B) Quantification of Rhesus macaque necropsy tissue DNA samples (from Rhesus macaque 27882) (including an uninfected negative control sample) for SIV DNA using ddPCR. (C) Comparison between ddPCR and nested qPCR analysis results for the necropsy tissue samples in (B).

In a set of tissue samples derived from a long-term cART suppressed Rhesus macaque (Figure 5B), we directly quantified SIV DNA viral load (total analyzed cell numbers range from 2.25 million to 10 million per sample) using the SIV ddPCR DNA assay on the Raindance platform and observed DNA viral loads ranging from 2 SIV DNA copies per million cells to 46 per million cells. The same set of samples (when replicate samples were available) was also analyzed using the qPCR approach where each sample was divided into 10 replicate reactions to minimize inhibition and allow the low-level viral quantity to be deduced from Poisson reaction positive rate. The ddPCR result in general agrees with the qPCR results, however we noticed that the ddPCR reads are 1.5 to 2 fold higher (Figure 5C) than the viral load obtained from the qPCR results, suggesting the possibility that there is still a moderate level of inhibition in the qPCR reactions (even when each sample was divided into 10 reactions).

### Establishing two-step RT-ddPCR reaction conditions for SIV RNA detection

In establishing a ddPCR protocol for detecting SIV RNA, we first attempted a one-step RT-ddPCR protocol. In this approach, reagents for a gene-specific reverse transcription priming step and PCR quantification step were combined with RNA template and subjected to dropletization. The one-step RT and PCR reactions were performed in droplets on a Bio-Rad C1000 Touch Thermal Cycler, followed by fluorescence detection on the RainDance Sense machine. This approach proved unsuccessful for low level SIV RNA detection (data not shown). We subsequently tested a two-step RT-ddPCR protocol, in which the reverse transcription step was performed prior to the formation of droplets. The resulting cDNA was then dropletized together with Taqman genotyping mastermix and the ddPCR assay primers and probe. PCR cycling of the droplets was performed on a Bio-Rad C1000 Touch Thermal Cycler, followed by fluorescence detection on the Sense machine. This approach was successful when gene specific priming (not random hexamer-priming) method was used for cDNA synthesis. In addition, the amount of reverse transcriptase greatly influenced the ddPCR detection outcome (Supplementary Table 3). The two-step RT-ddPCR assay can reliably detect single-digit copies of the SIV viral RNA (Figure 6A). The linear dynamic range of the assay for SIV RNA template was confirmed to be at least up to 1 million copies (upper limit tested) of viral nucleic acid per reaction (Supplementary Figure 3).

**Figure 6:**
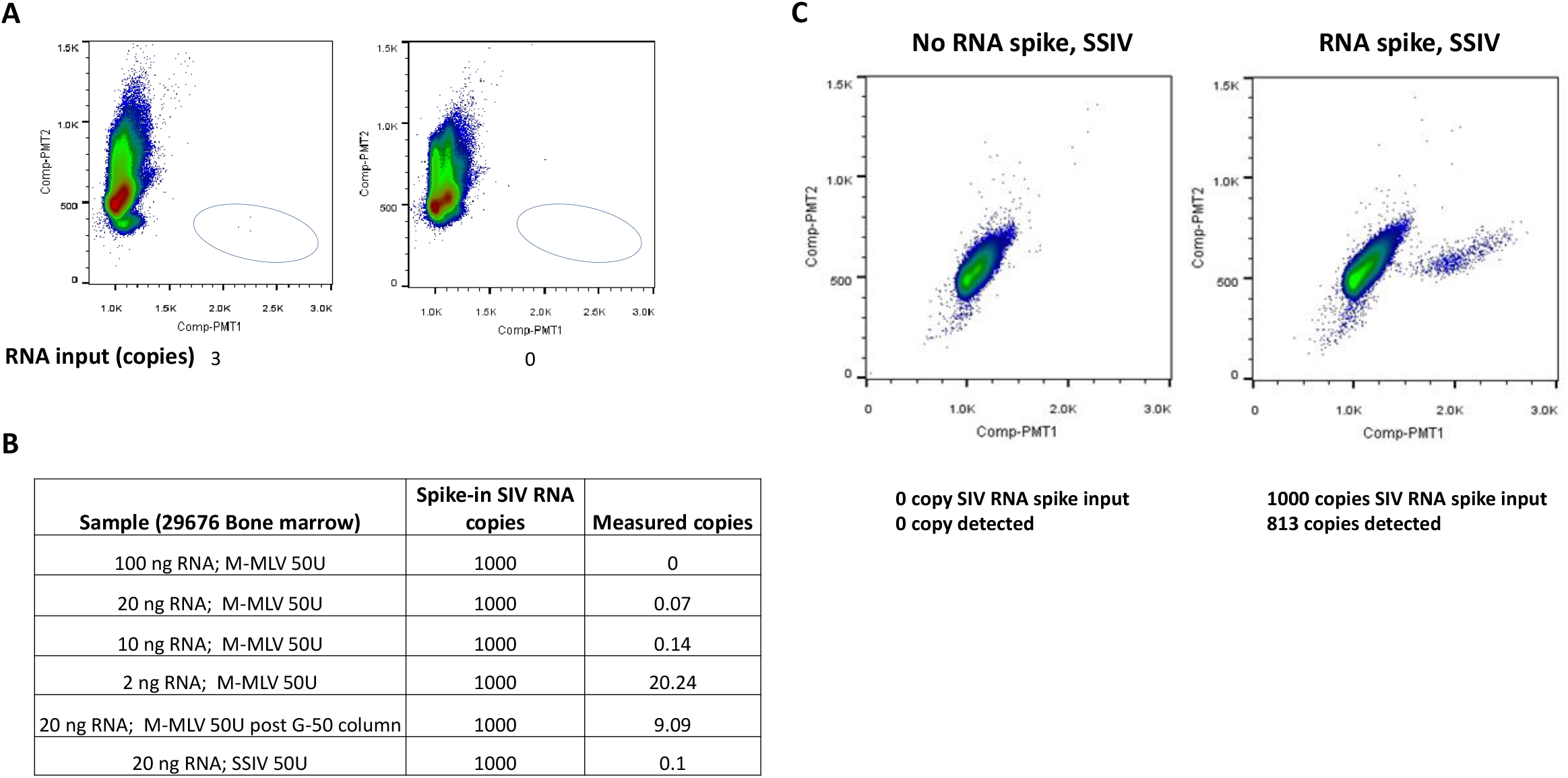
Superscript IV reverse transcription combined with Raindance ddPCR overcomes RNA sample inhibition. (A) Two step RT-ddPCR assay for SIV RNA detection. SIV RNA standard or buffer control were spiked in 1ug total RNA from naïve animal PBMC, and the samples were subject to reverse transcription (SIV gene-specific priming, RT enzyme: M-MLV, 200U/reaction), dropletization, end-point PCR and fluorescence reading/counting. (B) Severe RNA inhibition in a bone marrow aspirate sample (from Rhesus macaque 29676) in RT-qPCR analysis. Different amount of RNA sample extracted from a bone marrow aspirate sample, treated or untreated through a G-50 column as indicated, then subject to reverse transcription either using M-MLV or SSIV as the reverse transcriptase. The cDNA was subject to nested preamplification, and qPCR step was performed as described in Materials and Methods. The discrepancy between “measured SIV RNA copies” and “Spike-in SIV RNA copies” indicates the level of inhibition under each condition. (C) SSIV RT combined with SIV ddPCR assay overcomes bone marrow RNA sample inhibition. RNA was isolated from the bone marrow aspirate, and subject to SSIV reverse transcription (20 ng RNA per RT reaction). No preamplification step was performed on the cDNA.

### A high processivity reverse transcriptase combined with ddPCR overcomes RNA inhibition

Severe inhibition has previously been observed for certain Rhesus macaque tissue-derived RNA samples during the reverse transcription, PCR, or both steps. For example, samples 29676, 28808, 29211, 28885 and 29341 are RNA extracted from bone marrow needle aspirate samples from a long-term ART suppressed animal (i.e. expected RNA viral load is extremely low). Heparin, a known inhibitor of reverse transcription and PCR due to its high negative charge density and the ability to copurify with RNA (del Prete et al. 2007, Garcia et al 2002) was inadvertently introduced to anticoagulate the specimens during sample procurement. When 1000 SIV RNA copies were spiked into a reaction containing 100 ng 29676 RNA sample, no SIV RNA signal was detected by qPCR, indicating severe inhibition (Figure 6B). We explored different methods for potentially salvaging usable results from such heparinized specimens. Reducing the RNA sample input by 5 to 10-fold did not significantly alleviate the inhibition, while reducing the RNA sample input 50-fold allowed 20 copies (out of the 1000 copies) of input RNA to be detected (Figure 6B). Passing the RNA sample through a G-50 column slightly improved viral detection (Figure 6B), and subjecting the RNA samples to sequential precipitations with isopropanol and lithium chloride completely removed the inhibition (Supplementary Figure 4 left). However, the main observed disadvantage associated with Lithium chloride purification is that RNA recovery after lithium precipitation is often compromised, which is even more so when the sequential precipitation approach was used (Supplementary Figure 4 right). In an actual sample test scenario, this would effectively reduce the sample input (due to RNA loss during precipitation) for viral RNA analysis and reduce the test sensitivity. This would be less ideal especially in cases where the test samples are derived from animals receiving suppressive cART and have extremely low viral RNA load. In fact, the reduced effective sample input poses a detection sensitivity issue for any inhibitor removal method which cannot achieve consistent quantitative recovery.

As not all inhibitory factors introduced before and during RNA purification can be clearly identified, we sought to develop an approach that overcomes the inhibitor issue in RNA quantification that obviates the need for inhibitor removal. First, we searched for a reverse transcriptase that is less susceptible to inhibition. SuperScript IV (SSIV) is a high processivity reverse transcriptase that is capable of overcoming RT inhibition caused by biological or sample-prep inhibitors, such as isopropanol, ethanol, formalin, heparin, bile salts, guanidinium isothiocyanate, Trizol reagent, humic acid, among others. We noticed that SSIV-generated cDNA still showed almost complete inhibition when used as a template in qPCR quantification, consistent with heparin continuing to severely inhibit the PCR step (Figure 6B). When we combined SSIV generated cDNA with RainDance ddPCR quantification, 813 copies out of the 1000 copies of input RNA templates were detected (Figure 6C). Therefore, a 2-step RT-ddPCR procedure where SSIV enzyme was used in the reverse transcription step, and RainDance ddPCR was used in the PCR quantification step, could overcome severe RNA inhibition in bone marrow samples. This finding potentially expands the repertoire of tissue RNA samples that can be directly analyzed for viral RNA load without the need for removing inhibitors.

## Discussion

In this report, we describe the development and optimization of a SIV ddPCR DNA assay and a SIV RT-ddPCR RNA assay. These are ultrasensitive assays that, when performed on the RainDance digital droplet PCR platform, can detect single digit level DNA or RNA template quantities without a preamplification step. On a per reaction basis, the ddPCR DNA assay can tolerate significantly more DNA input compared to qPCR or Bio-Rad ddPCR platforms. In addition, we demonstrate that the combination of a high processivity RT enzyme and RainDance ddPCR could overcome inhibition in severely inhibited samples of extracted RNA. It is noteworthy that in the bone marrow RNA samples analyzed in this study, the likely dominant inhibitory factor was heparin. It will be of interest in the future to expand the study sample set to include inhibited RNA samples with unknown inhibitor identities.

We used a combination of SSIV in the reverse transcription step and RainDance ddPCR at the quantification step to overcome inhibition in SIV RNA measurements. Although the inhibitor (heparin) was not removed from the RNA sample and was present in both steps, the mechanisms for overcoming inhibition at the two steps were different. At the RT step, a high processivity enzyme, SSIV, was employed. By definition, a high processivity enzyme is a polymerase with a high probability to continue to copy the template rather than falling off, even in the presence of inhibitor molecules. Consistent with the importance of the reverse transcriptase’s processivity to overcome inhibition at the RT step, when Superscript III (SSIII), a lower processivity RT enzyme, was used in the RT step for the same severely inhibited bone marrow RNA sample, only 441 out of 1000 copies of the input RNA templates were detected (data not shown), which is about half of the detected signals of SSIV (441 copies vs. 813 copies). On the other hand, the ∼10 million droplets in the PCR step may contribute to segregating the inhibitor molecules and the cDNA template molecules into different droplets, i.e. reaction chambers, as these should be randomly distributed into droplets, which may explain an almost complete reversal of inhibition of RNA samples that were previously unanalyzable due to close to 100% inhibition during qPCR viral quantitation. In addition, endpoint detection of a digital readout is expected to be more tolerant of modest levels of reaction inhibition than real time methods.

The finding that RainDance ddPCR platform can tolerate significantly more input DNA than Bio-Rad ddPCR platform, even when equal reaction volumes were compared, was interesting. The average Raindance droplet (Raindrop) size is about 5 pL, compared to the claimed 1 nL volume of the Bio-Rad droplet size. The ∼200x difference in droplet size can be converted to ∼200x more droplets in a RainDance reaction compared to a Bio-Rad reaction with the same starting reaction volume, which can at least partly explain the significantly better ability of the RainDance platform to tolerate higher DNA input: more droplets could in principle allow better partitioning of inhibitory molecules from DNA template molecules into separate droplets (i.e. separate reaction chambers) to minimize the effects of potential inhibitors. Aside from potential inhibitors in DNA samples, excessive DNA input has previously been described to inhibit PCR through competing for polymerase binding (i.e. polymerase will bind to excessive input DNA rather than binding to the small quantity of duplex arising from primer binding to target site during the early cycles of PCR) (SantaLucia 2007). The dropletization step in ddPCR is expected to create a number of target sequence-containing droplets in which the effective ratio of the target sequence versus irrelevant DNA sequences is many magnitudes higher than such a ratio in a qPCR reaction. This should theoretically reduce competition for the DNA polymerase in target sequence-containing droplets, and increase the success rate of the PCR reaction in these droplets.

In conclusion, combining two ultrasensitive ddPCR assays for SIV nucleic acids detection with the Raindance ddPCR platform can enable significantly improved nucleic acid detection sensitivity by allowing a large amount of input DNA to be analyzed per reaction, and can overcome severe RNA inhibition when combined with suitable reverse transcription enzyme(s). These assays offer potential valuable tools for evaluating the treatment strategies aimed at reducing the latent reservoir and curing viral infection. The methodologies can also be adapted for additional applications where highly sensitive nucleic acid detection is required.

## Methods

### Nucleic acid extraction, qPCR quantification of proviral DNA and RT-qPCR quantification of tissue-derived SIV RNA

Nucleic acid extraction, preamplified quantitative real-time PCR quantification of proviral DNA and RT-qPCR quantification of tissue-derived SIV RNA followed procedures and conditions as described previously (Cline et. al 2005; Venneti et al 2008; Hansen et. al, 2013). Real time PCR step quantification was performed on ViiA 7 real time PCR system (Thermo Fisher Scientific).

For heparin removal from purified RNA samples, an isopropanol reprecipitation approach, a lithium chloride precipitation approach, or a sequential combination of the two approaches was used. For isopropanol reprecipitation, RNA pellet was first dissolved in 10 mM Tris pH8, followed by a 70% isopropanol precipitation and wash. For lithium chloride precipitation, the RNA pellet was first dissolved in 10 mM Tris 8.0, a 50% volume of 7.5M lithium chloride (Ambion AM9480) was added to the RNA solution, then a 2.5x volume of 100% ethanol was added to the mixture to allow precipitation of RNA followed by collection by microcentrifugation. RNA pellets were washed twice with 70% ethanol and air dried before being assayed.

### ddPCR optimization based on existing SIV gag assay format and master mix condition

We compared ddPCR to results obtained using an existing real time assay approach as detailed in Li et al, PNAS, 2016. For ddPCR modifications, the following reaction conditions were used: dNTPs (300 uM each), dUTP (600 uM), MgCl2 (varies; up to 5.5 mM), SGAG21 (forward primer) 600 nM, SGAG22 (reverse primer) 600 nM, pSGAG23 (FAM labeled probe, with a single BHQ quencher) 200 nM, 1x PCRII buffer (Thermofisher) with 0.2% Tween, AptaTaq (Roche) 1 unit, and 1x reaction stabilizer (RainDance).

Droplet formation was performed on the RainDrop Source instrument following the manufacturer’s instructions. During droplet formation, droplet integrity was monitored by visually examining a portion of the droplets in each lane as they moved through the device. In addition, total droplet count data for each sample after dropletization was retrieved from the Source instrument.

End point PCR was performed on a Bio-Rad C1000 Touch Thermal Cycler with the following PCR cycling condition: 95C 7min; 40 cycles of (95C, 15 sec; 60C, 1 min with a ramp rate of 0.5C per sec); 98C 10 min; 4C hold.

After thermocycling, droplet fluorescence reading was performed on the RainDrop Sense instrument per the manufacturer’s instruction. At the end of the run, total droplet count data for each sample after fluorescence reading was retrieved from the Sense instrument.

Data from Sense runs were analyzed using RainDrop Analyst to calculate the template copy number by modeling as a Poisson distribution. The formula used for Poisson modeling is:

Copies per droplet = -ln(1-p)

Where p = fraction of positive droplets.

For duplex assay Poisson modeling, the following definition and formula were used:

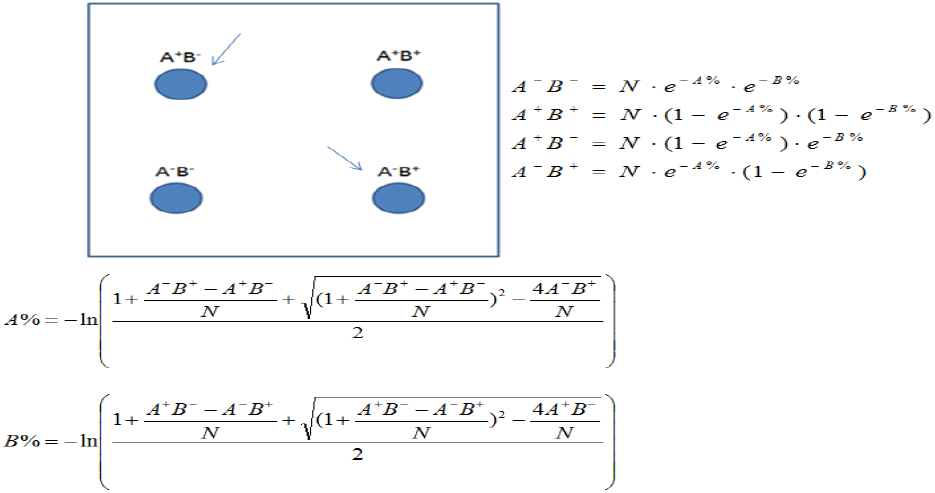

Where A-B-refers to drops containing neither target, A-B+ refers to drops containing target B only, A+B- refers to drops containing target A only, and A+B+ refers to drops containing both targets. N = the total number of droplet events.

### ddPCR quantification of proviral DNA

SIV Gag ddPCR primer sequences are as follows: SGag forward: GTCTGCGTCAT(dP)TGGTGCATTC; SGag reverse: CACTAG(dK)TGTCTCTGCACTAT(dP)TGTTTTG, whereas dP and dK (Lin & Brown, 1992) denote non-standard bases (Glen Research, Sterling, VA) introduced to minimize the impact of potential sequence mismatches at positions of described heterogeneity in SIV ioslates (Los Alamos Sequence Database, http://hiv-web.lanl.gov/). SGag ddPCR probe sequence is: 5’-FAM-CTT CYT CAG TRT GTT TCA CTT T-MGB. Rhesus macaque CCR5 ddPCR primer sequences are as follows: RCCR5 forward: CCAGAAGAGCTGCGACATCC; RCCR5 reverse: GTTAAGGCTTTTACTCATCTCAGAAGCTAAC and RCCR5 ddPCR probe sequence is. 5’ VIC-TTC CCC TAC AAG AAA CT-MGB.

Each 50 uL reaction mixture prior to dropletization included: 1x Taqman genotyping master mix (Thermofisher), SGag forward and reverse primers (600 nM each), SGag probe 200 nM, and/or RCCR5 forward and reverse primers (600 nM each), RCCR5 probe 200 nM, 1x reaction stabilizer (RainDance), appropriate amount of template (either in the form of SIV and/or CCR5 DNA standard, or DNA extracted from SIV infected or uninfected Rhesus macaque tissues). The concomitantly determined genomic CCR5 results were used to normalize viral DNA copy number results.

Droplet formation, end point PCR, Sense droplet reading, and data analysis were performed as described above.

### RT-ddPCR quantification of tissue-derived SIV RNA

A two-step RT-ddPCR approach was used for ddPCR quantification of tissue-derived RNA, as preliminary testing showed that a one-step RT-ddPCR approach did not perform well when both reverse transcription and ddPCR step reagents were incorporated into droplets before the one-step RT-ddPCR PCR cycling step. For the reverse transcription phase of the two-step protocol, each reaction contained 5 mM MgCl2, 500 nM of each dNTP, 1 mM DTT, 2 uM of SIVNestR01 (Hansen et al, 2011), 1x PCR II buffer (ThermoFisher) with 0.2% Tween, 10 U RNaseOUT, 200 U of M-MLV or SSIV (ThermoFisher) reverse transcriptase, and appropriate amount of SIV RNA standard, or tissue RNA from infected or uninfected Rhesus macaques, or molecular grade H2O. Before the droplet formation step, 15 uL of the reverse transcription product was combined with the following to yield a final reaction mixture: 1x TaqMan genotyping mastermix, SGag ddPCR forward and reverse primers (600 nM each), SGag ddPCR probe (200 nM), RCCR5 ddPCR forward and reverse primers (600 nM each), RCCR5 ddPCR probe (200 nM) and 1x reaction stabilizer (RainDance).

Dropletization, end-point PCR, Sense reading and data analysis were performed as described above.

### In vivo derived specimens

Specimens were graciously provided by Dr. Louis Picker (Oregon Health and Science University) and Dr. Paul Johnson (Emory University/Yerkes National Primate Research Center), from animals in protocols approved by the Institutional Animal Care and Use Committees at their respective institutions.

## Acknowledgements

The authors would like to acknowledge William Bosche for technical assistance, and Drs. Jeffrey Lifson and Robert Gorelick for helpful comments on the manuscript. We would also like to acknowledge Dr. Louis Picker (OHSU) and Dr. Paul Johnson (Emory University/YNPRC) for graciously providing in vivo derived specimens.

## Funding Disclosure

This project has been funded with Federal funds from the National Cancer Institute, National Institutes of Health, under Contract No. HHSN261200800001E. The content of this publication does not necessarily reflect the views or policies of the Department of Health and Human Services, nor does mention of trade names, commercial products, or organizations imply endorsement by the US Government.

## Competing Interests

The authors declare no competing interests.

## Supporting Information

Electronic supplementary information (ESI) available. See DOI:

## Supplementary Figure legends

**Figure S1. Probe and mastermix condition tests for SIV ddPCR DNA assay**. (A) Doubly quenched IDT probe test in SIV nested tissue DNA in the following master mixes: ACVP mastermix, Quanta ToughMix, TaqMan Universal mastermix, and TaqMan genotyping mastermix. (B) MGB and doubly quenched IDT probe test in SIV spike-in samples. The mastermix and probe combinations are as indicated. (C) Quanta genotyping mastermix test in cell DNA sample. (D) A summary of all conditions tested and the observed issue(s) associated with each condition.

**Figure S2. SIV ddPCR DNA assay performance**. (A) (B) Linear dynamic range of the SIV ddPCR DNA assay. Different amount of SIV DNA template was spiked in 1 million naïve cellequivalent of genomic DNA. SIV DNA amount was quantified by direct ddPCR. (C) Quantification data of the plots in (A) and Figure 4A.

**Figure S3. Two step RT-ddPCR assay for SIV RNA detection**. (A) Different copy numbers of SIV RNA standard were spiked in 1ug total RNA from naïve animal PBMC, and the samples were subject to reverse transcription (SIV gene-specific priming, RT enzyme: M-MLV, 200U/reaction), dropletization, end-point PCR and fluorescence reading/counting. (B) Linear dynamic range of the SIV RT-ddPCR RNA assay. (C) Quantification data of the plots in (A) and Figure 6(A).

**Figure S4. LiCl treatment of RNA sample removes heparin inhibition in RT-qPCR analysis but RNA recovery is low**. RNA samples from 4 bone marrow needle aspirate samples (from Rhesus macaque 28808, 29211, 28885 and 29341) were subject to additional precipitation step(s) (Isopropanol, lithium chloride, or a sequential combination of both) before SIV RNA quantitation analysis by adding 1000 copies of SIV RNA standard. Isopropanol or lithium chloride precipitation alone did not remove RNA inhibition, while isopropanol followed by lithium chloride precipitation completely removed heparin inhibition (left). However, lithium chloride precipitation often led to significantly and variably reduced RNA recovery.

**Table S1**. Total number of droplets generated during dropletization remains constant in the range of tissue DNA amount tested (1 million to 8 million cell equivalent input). Total droplet number for each sample after dropletization serves as an additional indicator of sample DNA input tolerance.

**Table S2**. MGB probe-based SIV ddPCR DNA assay characterization. Instead of measuring 60 ddPCR replicates to obtain 95% confidence, we obtained an approximate estimation of the LoD using a lower number of ddPCR reaction replicates and required the false negative rate to be below 5% (i.e. all 10 replicates have to be positive).

**Table S3**. Two step RT-ddPCR reverse transcription step condition tests. In these tests, SIV RNA standard was spiked in 1 ug cellular RNA isolated from naïve Rhesus macaque PBMC. Various conditions such as different reverse transcriptases (RT), RT enzyme amounts, and gene-specific vs. random hexamer priming methods were tested. The SIV RNA measured copies after ddPCR were compared to input copies to identify optimal reverse transcription conditions.

