## Supplemental figures for "Maximizing viral detection with SIV droplet digital PCR (ddPCR) assays"

### Figure S1A

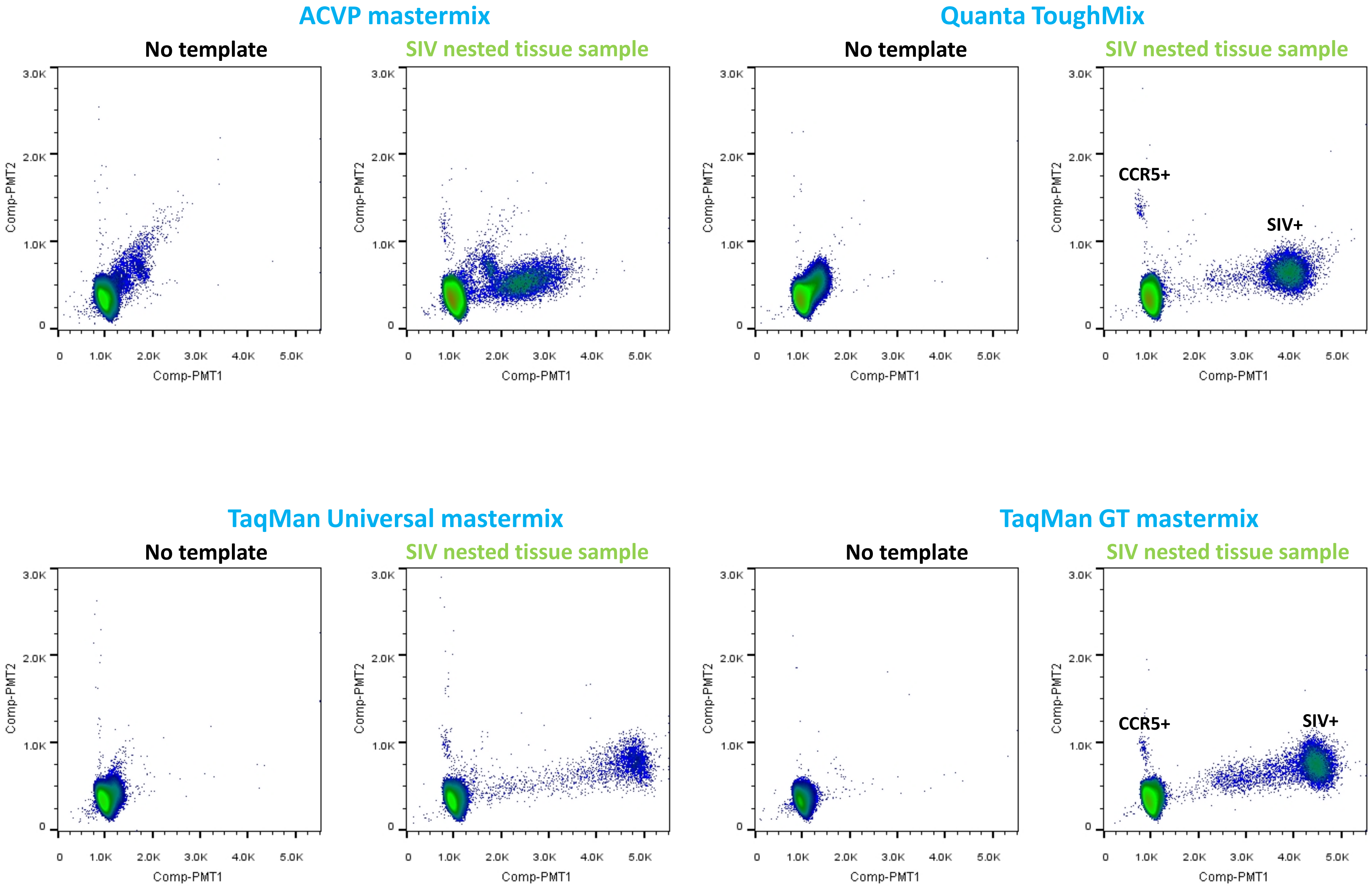


Doubly quenched IDT Zen probe used in all cases

#### Figure S1B

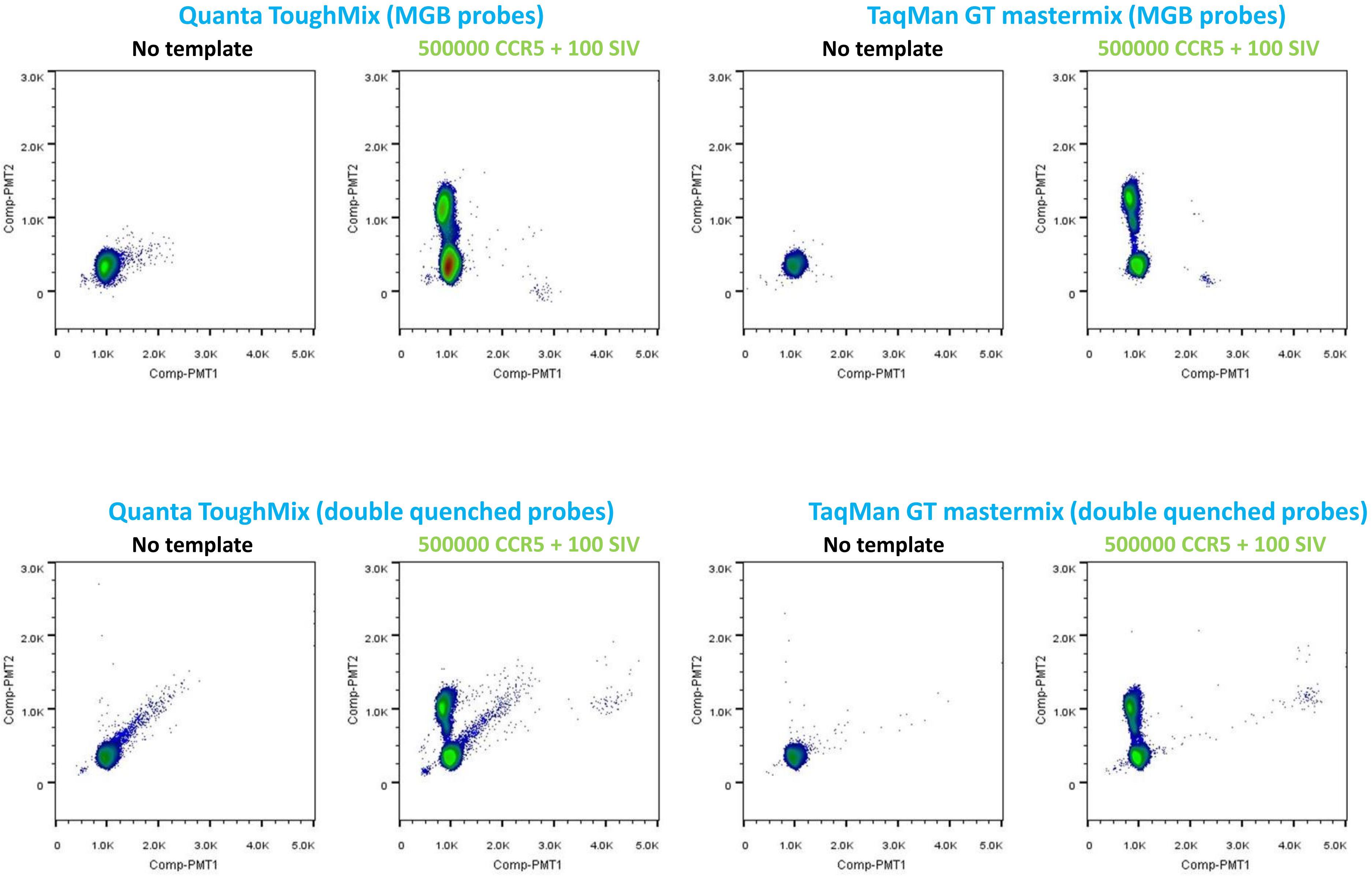

Quanta ToughMix (MGB probes)
TaqMan GT mastermix (MGB probes)
Quanta ToughMix (double quenched probes)
TaqMan GT mastermix (double quenched probes)
No template
500000 CCR5 + 100 SIV
Comp-PMT1
Comp-PMT2
0
1.0K
2.0K
3.0K
4.0K
5.0K

### Figure S1C

Quanta Genotyping MasterMix used in all cases

**No template, double quenched probes**

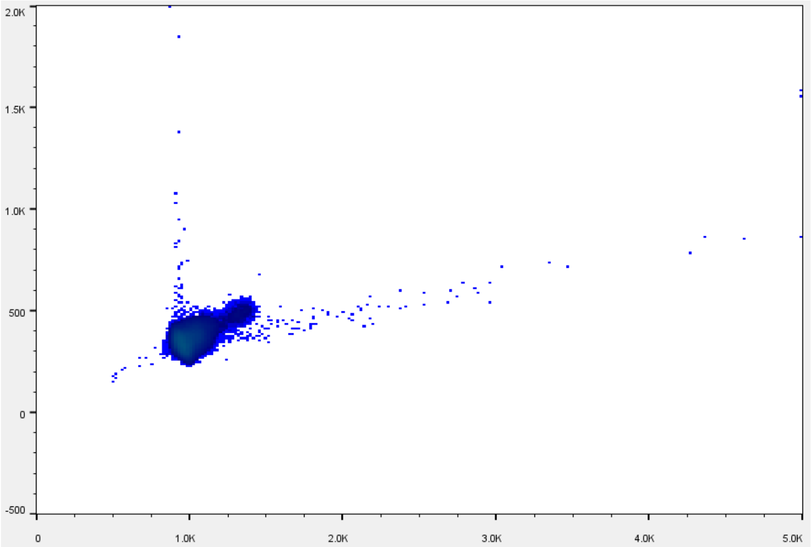


**1M cell input, double quenched probes**

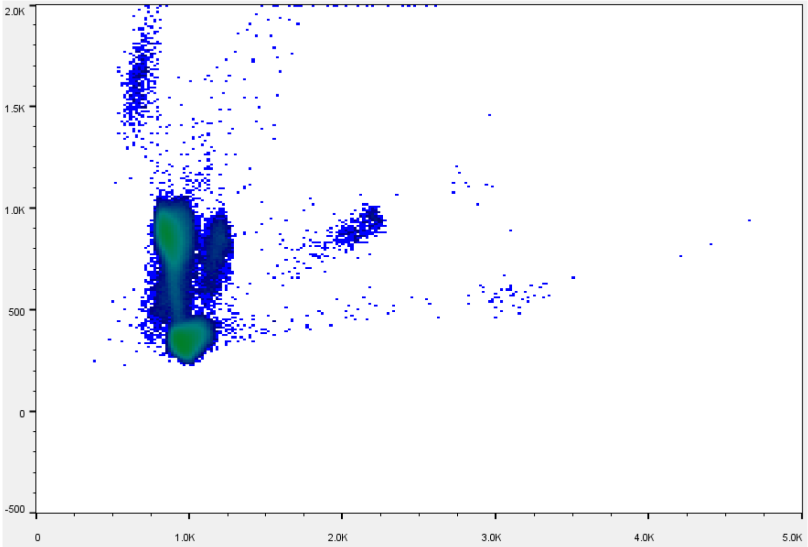


**No template, MGB probes**

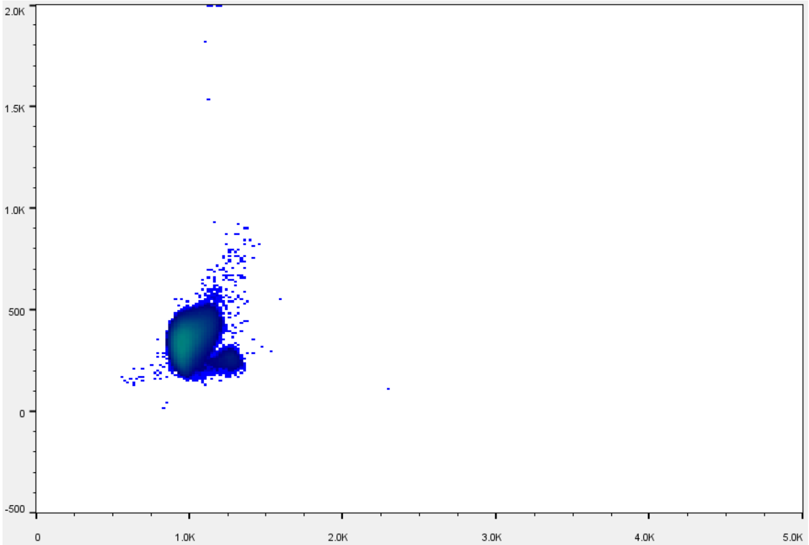


**1M cell input, MGB probes**

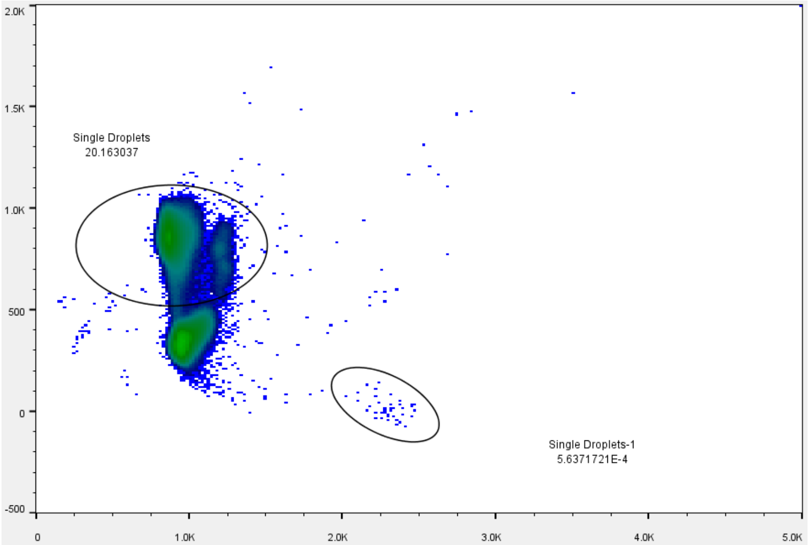

#### Figure S1D

| Figure | Probe | Template | Mastermix | Issue(s) |
|---|---|---|---|---|
| Sup 1A | Double quenched | SIV nested tissue DNA | ACVP | Cluster separation; low background (in no template reaction) |
| Sup 1A | Double quenched | SIV nested tissue DNA | Quanta ToughMix | Low background |
| Sup 1A | Double quenched | SIV nested tissue DNA | TaqMan Universal | Low count (2842 out of 10000); low background |
| Sup 1A | Double quenched | SIV nested tissue DNA | TaqMan Genotyping | Low background |
| Sup 1B | MGB | Spiked-in CCR5 and SIV | Quanta ToughMix | Cluster tightness |
| Sup 1B | Double quenched | Spiked-in CCR5 and SIV | Quanta ToughMix | Cluster tightness |
| Sup 1B | Double quenched | Spiked-in CCR5 and SIV | TaqMan Genotyping | Cluster tightness; low background |
| 2A | MGB | Spiked-in CCR5 and SIV | TaqMan Genotyping | |
| Sup 1C | Double quenched | Ovarian tissue DNA | Quanta Genotyping | Diffuse cluster; low background |
| Sup 1C | MGB | Ovarian tissue DNA | Quanta Genotyping | Diffuse cluster |

#### Figure S2A

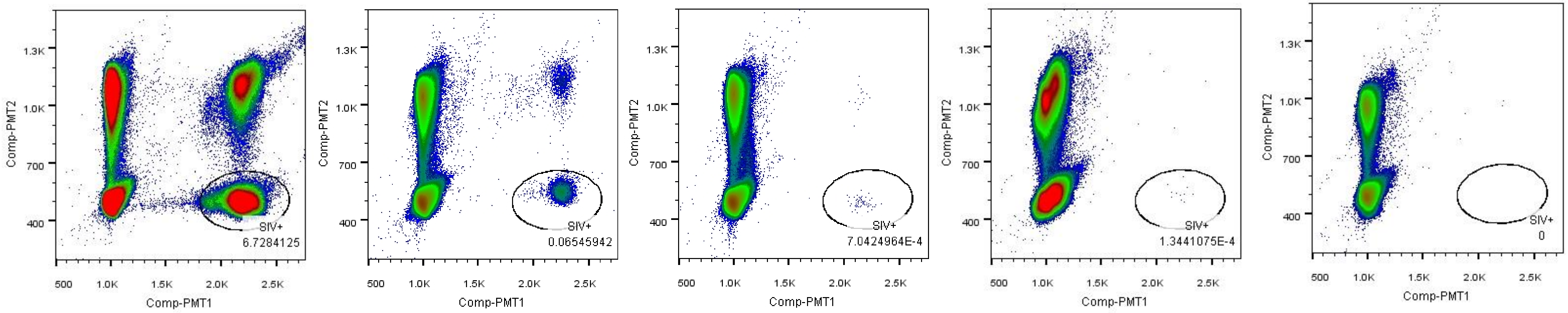


#### Figure S2B

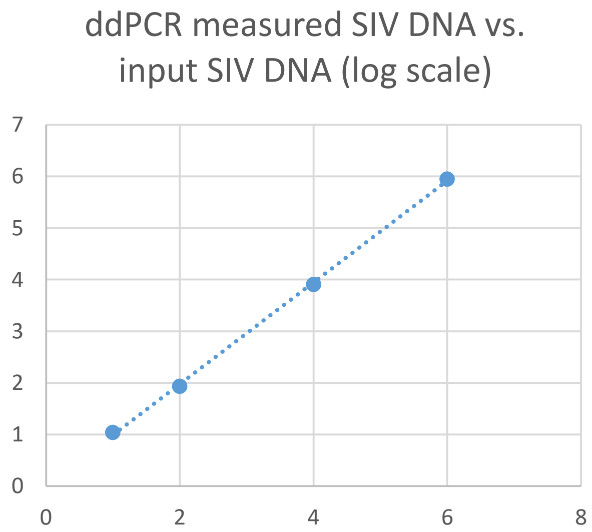


#### Figure S2C

| Cell count (CCR5) | SIV DNA standard input | SIV DNA count |
|---|---|---|
| 1162866 | 1 million | 888036 |
| 1074749 | 10000 | 8135 |
| 990689 | 100 | 86 |
| 1087174 | 10 | 11 |
| 1115984 | 3 | 5 |
| 1247780 | 2 | 2 |
| 1184228 | 0 | 0 |

#### Figure S3A

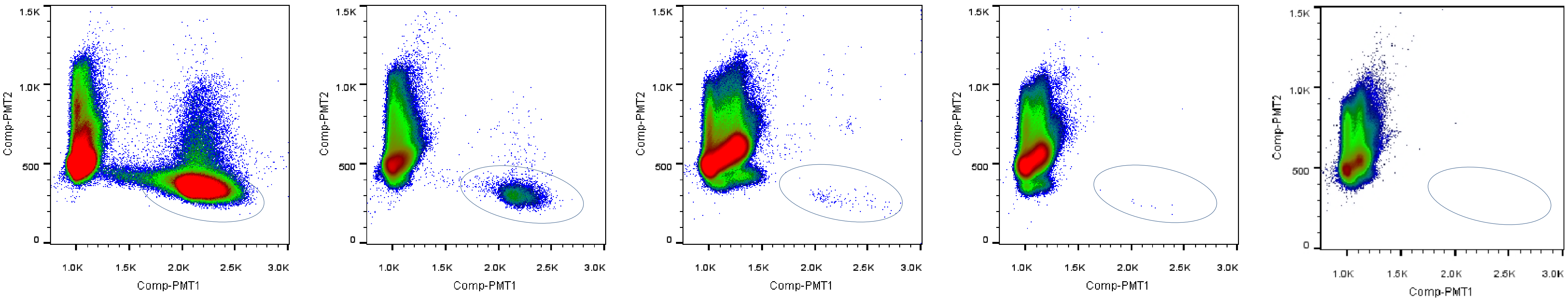


**RNA input (c)** 1 million 10,000 100 10 0

#### Figure S3B

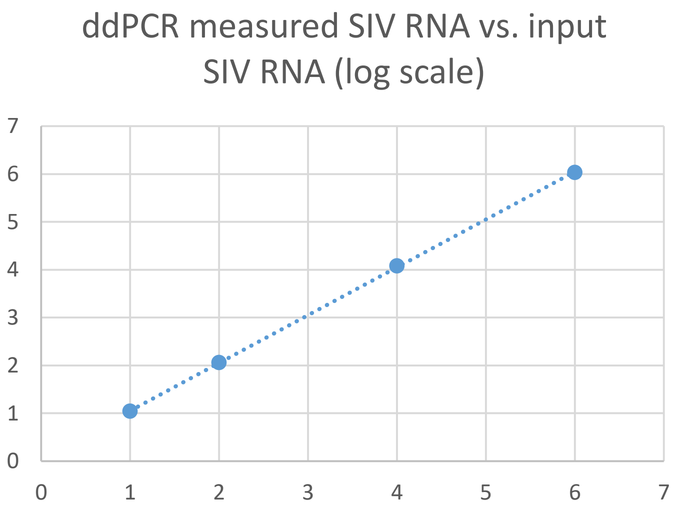


#### Figure S3C

| SIV RNA standard input | SIV RNA measured |
|---|---|
| 1 million | 1076191 |
| 10000 | 11922 |
| 100 | 115 |
| 10 | 11 |
| 3 | 3 |
| 0 | 0 |

### Figure S4

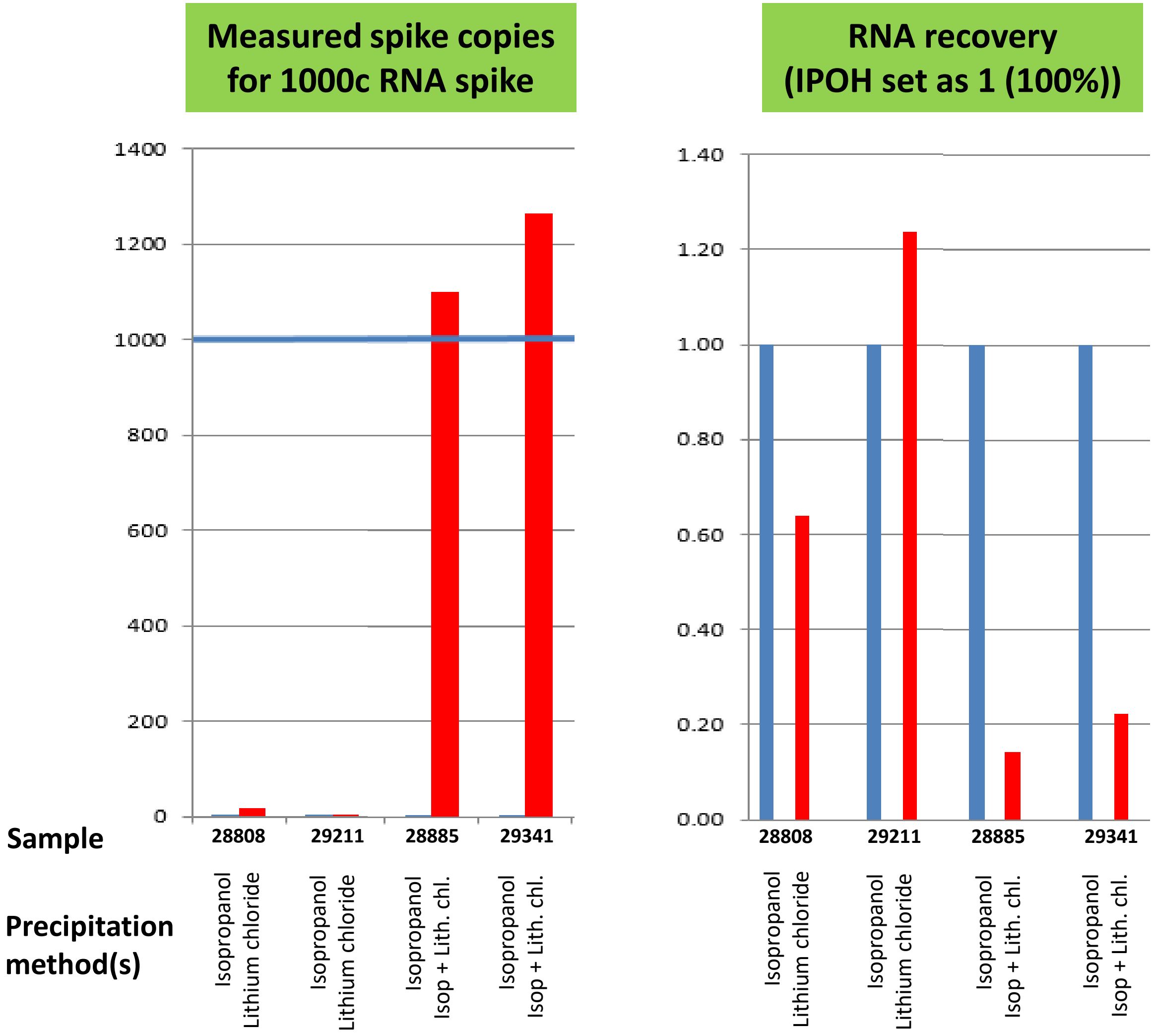
Measured spike copies for 1000c RNA spike
RNA recovery (IPOH set as 1 (100%))
1400
1200
1000
800
600
400
200
0
1.40
1.20
1.00
0.80
0.60
0.40
0.20
0.00
Sample
28808
29211
28885
29341
Precipitation method(s)
Isopropanol
Lithium chloride
Isopropanol
Lithium chloride
Isopropanol
Isop + Lith. chl.
Isopropanol
Isop + Lith. chl.

#### Table S1

| Cell equivalent | Droplets |
|---|---|
| 1 million | 164243 |
| 2 million | 164426 |
| 3 million | 160534 |
| 4 million | 159180 |
| 5 million | 154094 |
| 6 million | 155404 |
| 7 million | 158426 |
| 8 million | 162222 |

### Table S2

| Reaction | A | B | C | D | E | F | G | H | I | J | K |
|---|---|---|---|---|---|---|---|---|---|---|---|
| **Cell input** | 1M | 1M | 1M | 1M | 1M | 1M | 1M | 1M | 1M | 1M | 1M |
| **SIV DNA input** | **3** | **3** | **3** | **3** | **3** | **3** | **3** | **3** | **3** | **3** | **0** |
| **SIV DNA count** | **3** | **2** | **3** | **3** | **6** | **3** | **3** | **6** | **1** | **5** | **0** |

**Table S3**

| RT enzyme | Amount | Priming | SIV RNA input (copies) | SIV RNA measured (copies) |
|---|---|---|---|---|
| **M-MLV** | 200U | GSP | 10 | 14 |
| **M-MLV** | 50U | GSP | 10 | 5 |
| **SSIII** | 200U | GSP | 10 | 14 |
| **SSIII** | 20U | GSP | 10 | 3 |
| **M-MLV** | 200U | HEX | 10 | 1 |
| **SSIII** | 200U | HEX | 10 | 2 |
| **SSIII** | 20U | HEX | 83 | 35 |
| **SSIII** | 200U | HEX | 83 | 84 |
| **SSIII** | 20U | GSP | 83 | 74 |
